# Regulation of a Classical Allosteric Molecular Machine by an Intrinsically Disordered Domain: the C-termini of GroEL

**DOI:** 10.64898/2026.09.01.748644

**Authors:** Morgan E. Powers, Kacie A. Evans, He (Mirabel) Sun, David H. Russell, Hays S. Rye

## Abstract

The bacterial chaperonin GroEL is a canonical example of an ATP-dependent molecular machine that must couple ligand binding to productive conformational work. GroEL passes through a series of distinct structural shifts, driven by ATP binding and hydrolysis, which power a facilitated protein folding reaction. How the complex allostery of the GroEL oligomer creates a folding cycle that is both efficient and directional remains incompletely understood. Here, we combine variable-temperature native ion mass spectrometry with single-molecule FRET to examine how the intrinsically disordered, highly conserved GroEL C-terminal tails impact the allosteric behavior of a single GroEL ring. Our observations show that the C-terminal tails restrain the conformational dynamics of the GroEL ring, most likely through direct interactions with the upper apical domains of the GroEL subunits, a constraint that is progressively released as ATP binds. These results support a model in which the C-terminal tails act as an entropic regulator of the GroEL reaction cycle: transient interactions between the tails and GroEL apical domains restrain premature ring opening and tune the energetic threshold for productive engagement by the smaller GroES co-chaperonin. By linking disordered tail dynamics to the classically cooperative reorganization of the GroEL ring, this mechanism enforces an ordered allosteric cascade that minimizes wasteful formation of empty GroEL-GroES cavities. These findings reveal how the conformational properties of an intrinsically disordered element can be exploited to optimize the energetic efficiency and functional timing of a large allosteric machine.

**SIGNIFICANCE:** How large molecular machines avoid wasteful ATP consumption while maintaining productive reaction cycles remains a central question in mechanistic biology. Here, we identify a regulatory role for the conserved, intrinsically disordered C-terminal tails of GroEL, one of the best-studied ATP-dependent chaperones. Our data indicate that transient interactions between the tails and GroEL apical domains restrain premature ring opening and tune the energetic threshold for productive binding of the smaller GroES co-chaperonin. By linking disordered tail dynamics to the cooperative reorganization of the GroEL ring, this mechanism enforces an ordered allosteric cascade that minimizes wasteful formation of empty GroEL-GroES cavities. These findings illustrate how intrinsically disordered regions can function as tunable regulatory elements that optimize the efficiency of complex biological machines.

## INTRODUCTION

One of the great challenges faced by all living organisms is the need to maintain a healthy proteome. Production of active proteins must occur at rates sufficient to maintain life but under conditions that unavoidably enhance protein folding errors like misfolding and aggregation (1, 2). Complex folding topologies combined with the concentrated and crowded conditions of a living cytoplasm present enormous obstacles to efficient folding (3–5). Unpredictable external environmental shifts, like thermal stress, exacerbate the severity of the problem (6, 7). The need for proteome surveillance and maintenance, generally referred to as proteostasis, imposes a large energetic load on all organisms (8, 9). Because the evolutionary race for survival creates an unforgiving fitness constraint, such expenditures of precious metabolic energy must be managed with minimal waste.

Proteostasis is built upon a foundation of energy-utilizing, cellular nano-machines, which monitor, fold, fix and, eventually, clear proteins that are either damaged or that have simply outlived their usefulness (8, 10). Constructed from several different families of molecular chaperones, folding sensors and regulated proteases, the proteostasis network ensures efficient production of active proteins while clearing the folding and assembly errors that result in protein aggregation and toxicity. The Hsp60s or chaperonins, one of the first families of molecular chaperones to evolve, occupy a central node of this network (10, 11). These large, oligomeric cellular machines employ ATP hydrolysis to facilitate the folding of a subset of proteins that are exceptionally poor at, or even incapable of, spontaneous folding (12, 13). In general, this catalyzed folding process takes the form of a cyclic substrate protein capture, confinement and release sequence (14–16). The canonical bacterial Hsp60, known as GroEL, is constructed from two, seven-membered rings that are stacked back-to-back to form a tetradecamer complex with two, solvent filled open cavities (17). Capture of a kinetically trapped folding intermediate on an open GroEL ring is followed by ATP binding, which permits GroES (a smaller, co-chaperonin heptamer) to bind (18–20). Stable GroES binding causes displacement of the folding intermediate into a closed, privileged cavity inside the GroEL-GroES complex where folding is initiated (19–21). Hydrolysis of ATP inside this complex, followed by a new round of substrate protein and ATP binding to the second GroEL ring, triggers disassembly of the folding cavity, release of GroES and ejection of the previously confined folding intermediate back into free solution (22–25).

At its core, the GroEL folding cycle depends on a complex allosteric interplay between conformational states of the GroEL subunits and ATP. Early studies demonstrated that GroEL behaves like a sophisticated, but classical, allosteric machine with positively cooperative binding of ATP to one ring, followed by negatively cooperative (i.e. inhibitory) ATP binding to the second (26). This well-accepted nested cooperativity model is supported by a large body of biochemical and structural data collected over many years (26–31). However, the physical basis of allostery has been dramatically revised over the same period of time (32–37). From deterministic frameworks built around mechanical analogies (e.g. pulleys, levers and springs), these reformulations emphasize a probabilistic view of allostery, where ligand binding shifts the micro-state distribution of a system over a range of potential length and time scales (34, 38). While these redistributions can be hidden at the ensemble level, allosteric responses are ultimately grounded in which subset of micro-states are populated and for how long. This modern dynamic framing of allostery can, for example, explain how intrinsically disordered proteins, which defy classical description, possess allosteric responses (39–41). Importantly, several studies have shown that GroEL displays a greater level of micro-state heterogeneity than originally recognized, suggesting that the allosteric mechanism of this chaperonin, at least as outlined by classical models, is likely incomplete (42–45).

One dynamic and poorly understood feature of the GroEL oligomer is the intrinsically disordered carboxy-terminal extension that protrudes into the central cavity from the inner, bottom face of each subunit. These approximately 2 kDa, amphipathic tails are enriched in GGM repeats and are conserved in the sub-family of Hsp60s to which GroEL belongs (the so-called Type I chaperonins) (46). While an early study found that the tails can be deleted without compromising tetradecamer formation, stability or bacterial survival under laboratory conditions, tail removal was associated with a substantial, though mysterious, fitness cost (47). More recent work has shown that alterations to, or removal of, the tails can change the kinetics of ATP hydrolysis and reduces, but does not eliminate, facilitated folding of several substrate protein models *in vitro* (48–54). We previously demonstrated that the classical, ATP-driven allosteric response of a tailless GroEL tetradecamer (ELΔ526) is profoundly altered (50). We and others have also shown that the C-terminal tails make direct physical contact with non-native folding intermediates, both on an open GroEL ring and after encapsulation beneath GroES (49, 51, 55, 56). Additionally, we demonstrated that the tails are involved in substrate protein retention and forced unfolding during assembly of the GroEL-GroES complex (49–51). In total, these observations suggest that the intrinsically disordered tails are coupled to the allosteric cycle of the GroEL oligomer in a functionally relevant way. The nature of this connection, and how the conformational states of well-structured GroEL domains could be linked to highly dynamic micro-states of the disordered tails is unclear.

Isolating which micro-states of the GroEL oligomer map to functional, macro-state outputs, and determining how the intrinsically disordered C-termini impact these distributions, is challenging. Traditional biophysical measurements smear these microscopic distributions into ensemble averages across both the population of tetradecamers and the fourteen subunits of each oligomer. A recent smFRET study successfully reduced the severity of this averaging problem by creating a variant of the well-studied single ring variant of GroEL (SR1) carrying one fluorescently labeled subunit (42). This study demonstrated that not only does SR1 possess significant micro-state dynamics, but the observed distribution shifts to more open micro-states in response to ATP, paralleling the classical macro-state behavior of a GroEL ring. However, this study permitted only limited insight into how ATP ligation state is linked to shifts in micro-state dynamics of the SR1 ring (42).

We and others have shown that native ion mass spectrometry (nMS) using modern ultra-high mass detectors can resolve the ligation state distributions of the GroEL and SR1 oligomers in the presence of ATP and ADP (57, 58). Combined with a novel approach to sample temperature control, referred to as variable temperature nMS (vT-nMS), we have shown that the site-resolved thermodynamics of nucleotide binding to a GroEL ring can be quantified (58–62). Here, we combine vT-nMS and smFRET to examine the role of the intrinsically disordered C-terminal tails in the response of the GroEL ring to ATP. Using SR1 as a model, our results suggest that the tails function as an “entropic latch,” in which direct physical contact between the tails and the GroEL apical domains acts to damp the conformational dynamics of both the tails and apical domains. As ATP binding pushes the ring toward more open micro-states, these tail-apical restraints are released, providing a large entropic contribution to the progression of microscopic ATP binding free energies. We propose that the entropic tuning provided by the tails functions as a central allosteric integrator for the GroEL reaction cycle, optimizing the formation of substrate protein-occupied GroEL-GroES folding cavities while minimizing the energetically wasteful formation of empty complexes.

## RESULTS

### Deletion of the C-terminal tails increases both ATP binding affinity and cooperativity

To examine how the C-terminal tails impact the allosteric mechanism of GroEL, we employed native ion mass spectrometry (nMS) to quantify ATP binding to the well-studied single ring GroEL variant (SR1) (19, 21), in the presence and absence (SRΔ526) of the tails (49, 50). Uniquely, nMS permits the binding of ATP to individual SR1 subunits to be measured as a function of ATP concentration. Both SR1 and SRΔ526 were first exchanged into a charge-reducing, mass spectrometry-compatible buffer system (200 mM EDDA, pH = 6.3, 1 mM MgOAc), in which ATP binding to GroEL subunits can occur but hydrolysis cannot. Removal of the C-terminal tails results in a substantial decrease in the ATP concentration needed to fully saturate the heptamer ring (Figure 1A and B; SR1 versus SRΔ526). A plot of the fractional saturation of fully liganded SR1 rings (i.e., n = 7 bound ATPs per oligomer) as a function of ATP concentration, yields a sigmoidal plot that is well fit by Hill’s equation (Figure 1C) with a Hill constant (n_H_) of 2.6, in excellent agreement with prior bulk measurements of ATP binding to SR1 (26, 57, 63, 64). Removal of the C-terminal tails results in a large increase in observed ATP binding cooperativity for SRΔ526 (n_H_ = 3.2) and a decrease in the half saturation concentration from 32 µM for SR1 to 8 µM for SRΔ526.

**Figure 1.**
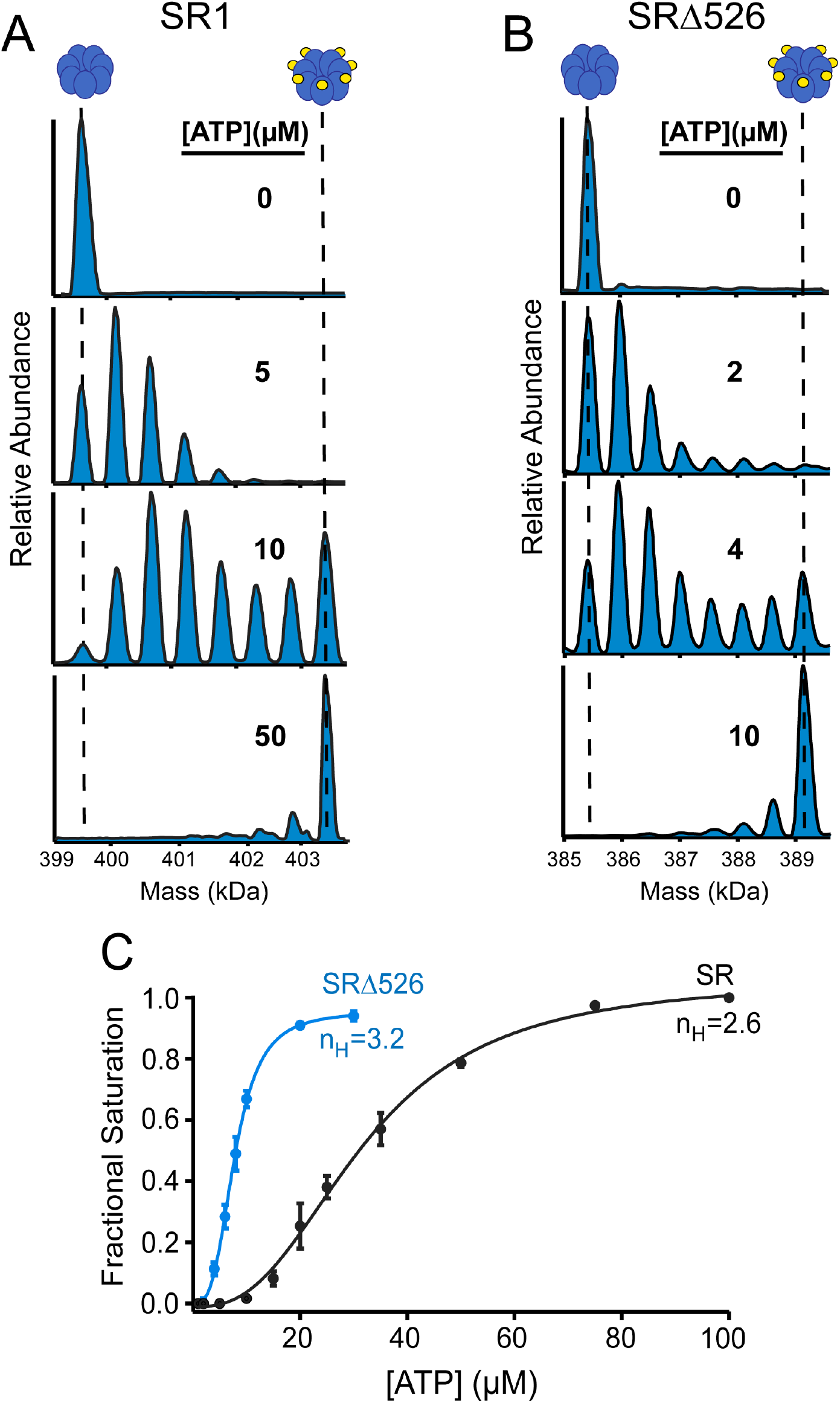
Deletion of the intrinsically disordered C-terminal tails enhances the cooperativity of ATP binding to a GroEL ring. Deconvolved native ion mass spectra of (A) 1 µM single-ring GroEL (SR1) and (B) 1 µM SRΔ526 (a C-terminal SR1 deletion mutant) in 200 mM EDDA, 1 mM Mg(OAc)_2_ at 20 °C mixed with different concentrations of ATP. Each ligation state of the SR1 and SRΔ526 oligomers can be resolved as a distinct mass peak, ranging from n = 0 (apo) to n = 7 (fully saturated) bound ATP. SR1 and GroEL do not appreciably hydrolyze ATP under these experimental conditions (62). (C) Plot of the fractional saturation of the fully liganded (n = 7) rings for SR1 and SRΔ526 as a function of total ATP concentration. Fractional saturation curves were fit to the Hill equation resulting in a n_H_ for SR1 of 2.6 ± 0.2 and half saturation of 32 ± 1.2 µM and a n_H_ for SRΔ526 of 3.2 ± 0.2 and half saturation of 7.8 ± 0.1 µM.

We next explored how the affinity of each nucleotide binding site, as a function of oligomer saturation, is affected by the presence of the C-terminal tails. We applied a variable temperature enhancement to our nMS approach (vt-ESI-nMS) previously employed to study nucleotide binding to GroEL and SR1 (58–62). Consistent with these prior studies, we found that ATP binding to both SR1 and SRΔ526 is strongly temperature dependent (Figure 2A and B). Removal of the C-terminal tails results in a dramatic shift of the SRΔ526 ring toward the fully saturated state of the oligomer at all temperatures (Figure 2B). Apparent ATP binding constants (K_a_) for individual subunits of the SR1 and SRΔ526 oligomers were extracted from fits to a macroscopic sequential binding model (Figure 2 C and D) (65). While both SR1 and SRΔ526 demonstrate a progressive increase in site affinity as ring saturation increases, as expected for cooperative nucleotide binding within a GroEL ring, individual site affinities for SRΔ526 are between five and ten-fold higher than those observed for SR1. Additionally, while ATP binding to both SR1 and SRΔ526 displays a strong dependence on temperature, the SR1 ring with its intact C-termini appears to be much more sensitive to shifts in temperature, with a considerably larger reduction in ATP binding as the temperature increases (Figure 2A and B).

**Figure 2.**
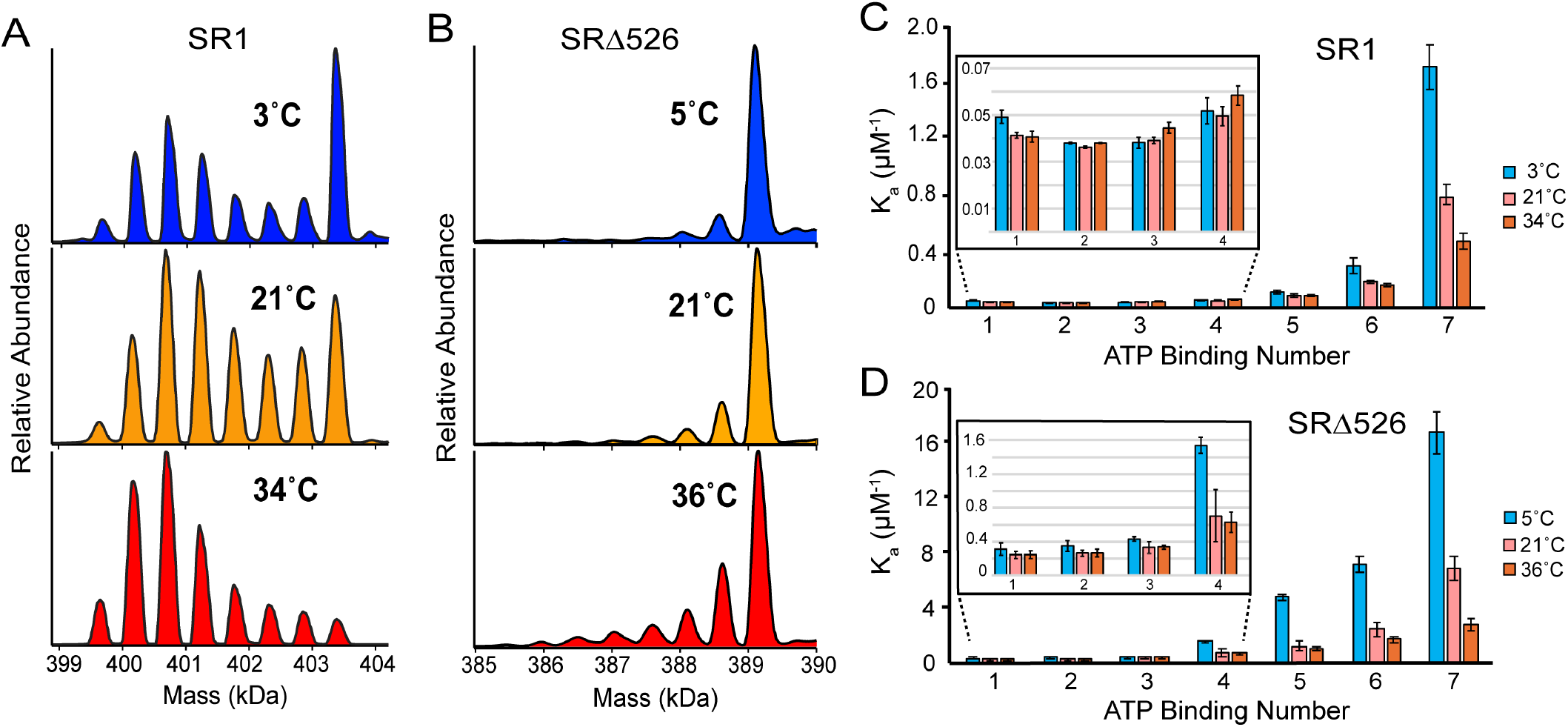
C-terminal tail deletion increases the ATP binding affinity of each SR subunit by approximately 10-fold. Deconvolved mass spectra of (A) 1 µM SR1 and (B) 1 µM SRΔ526 in 200 mM EDDA, 1 mM Mg(OAc)_2_ and 10 µM ATP at various temperatures. Increasing temperature reduces ATP binding affinity in all cases. Apparent ATP binding constants (K_a_) for individual subunits of the (C) SR1 and (D) SRΔ526 oligomers were extracted from fits to a sequential macro-state binding model (65).

### Removal of the C-terminal tails changes the thermodynamics of ATP binding

By extracting the temperature-dependent, per site nucleotide binding free energies from data collected at different ATP concentrations (Figure S1), it is possible to determine the enthalpy and entropy changes associated with each individual ATP binding event using van’t Hoff analysis (60–62, 66). As expected for SR1, the observed binding free energy for the first three binding events are all approximately the same (Figure 3A). However, the 4th ATP binds with a slightly higher free energy (ΔΔG° ∼ 0.6 kJ/mol), while the 5th through the 7th binding events display progressively larger binding free energies (ΔΔG° 1.2 - 3.3 kJ/mol per binding event). This behavior is consistent with the expected macro-state transition of the SR1 ring from a lower affinity T-like state to a higher affinity R-like state once the ring has bound 3-4 ATPs. Strikingly, the binding of the first six ATPs is dominated by a large and favorable entropy change, with a negligible enthalpic contribution until the 6th ATP molecule binds (Figure 3C). By contrast, removal of the C-terminal tails results in a dramatic shift in the enthalpic and entropic contributions to ATP binding (Figure 3D). While the first three ATP binding events to SRΔ526 remain entropy-dominated, enthalpy plays a larger role than is seen with SR1. By the 4th binding event, the free energy of ATP binding is equally partitioned between favorable enthalpic and entropic contributions for SRΔ526. Distinct from SR1, subsequent ATP binding is almost completely enthalpically dominated, with entropy contributing little to the free energy change of the 5th to 7th ATP binding events for SRΔ526. These observations strongly suggest that the allosteric design of the GroEL ring rests, in part, on the ATP-driven release of a large and energetically unfavorable conformational and/or hydration state of the T-state subunits. These results also show that the dynamic, intrinsically disordered C-terminal tails play a central role in this process.

**Figure 3.**
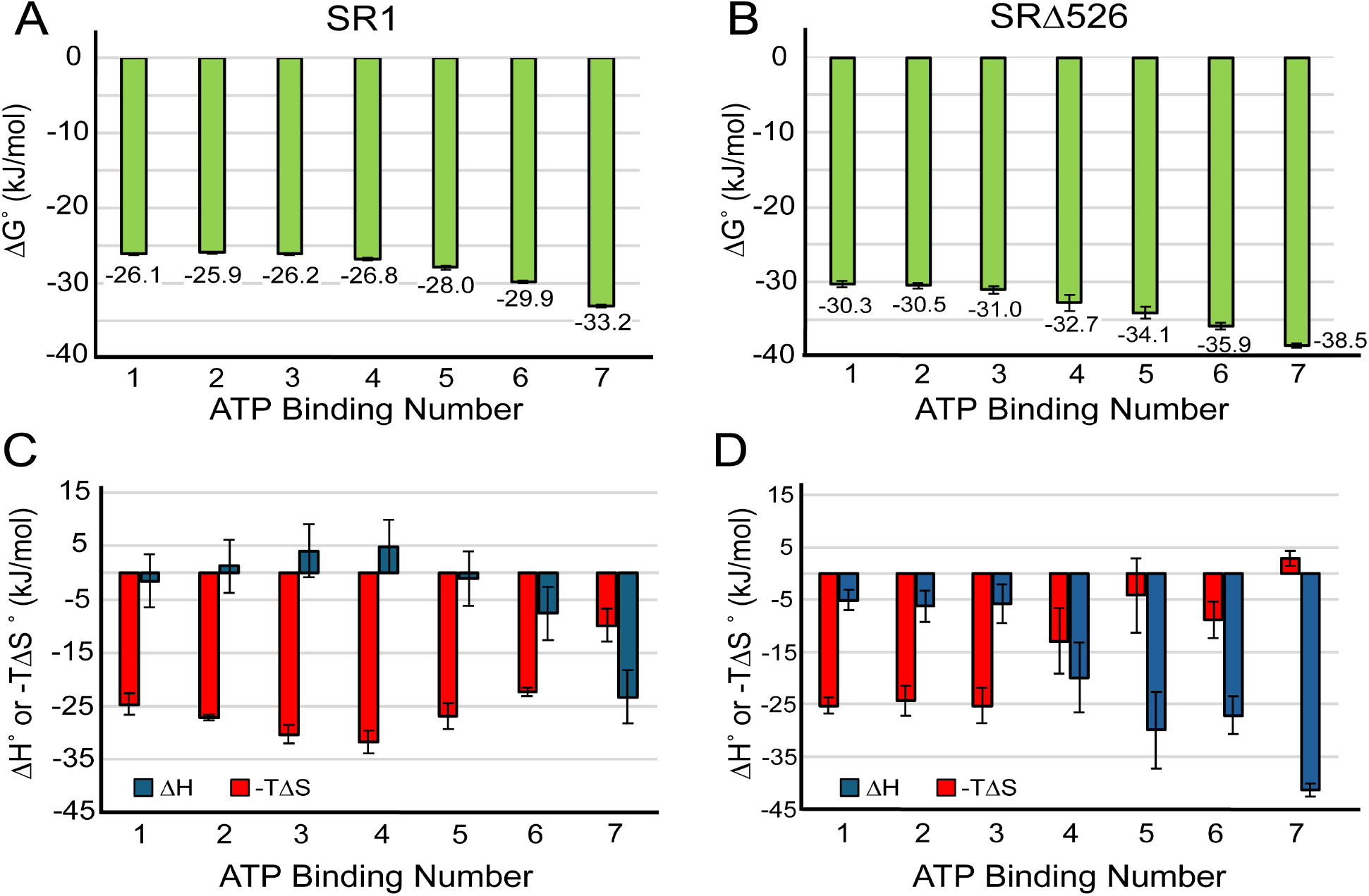
The thermodynamic signature of ATP binding to SRΔ526 is distinct from that of SR1. The standard state free energy difference (ΔG°) associated with each ATP binding event for (A) SR1 and (B) SRΔ526 at 20 °C was calculated from the observed macro-state binding constants for each ligation state. The changes in standard state enthalpy (ΔH°) and entropy (ΔS°) for each ATP binding event were calculated for (C) SR1 and (D) SRΔ526 by fitting the observed temperature dependence of the site-resolved binding free energies to a non-linear version of the van’t Hoff equation (61, 65). ATP binding to the first six SR1 subunits is driven almost exclusively by a large, favorable change in entropy (-TΔS°). For SRΔ526, only the first three ATP binding events are entropy dominated while the free energy change of the last three to four ATP binding events is dominated by a favorable enthalpy change.

### The C-terminal tails alter the microstate distributions and interconversion kinetics of the apical domains

To understand how the C-termini influence the conformational dynamics of SR1, we employed a previously described single molecule Förster resonance energy transfer (smFRET) assay that captures changes in the micro-state distribution of the ring as it binds nucleotides (Figure 4A) (42). In the absence of ATP, SR1 appears to populate two resolvable micro-states (S_1_, S_2_) with respect to elevation of the apical domains (Figure 4B). Micro-states were identified using a minimal multi-Gaussian peak fit of the observed smFRET efficiency histograms (Figure S2). Addition of low ATP concentrations (1 µM) results in appearance of a third, lower FRET state (S_3_), likely corresponding to the most open, R-like conformation of the subunit. As the ATP concentration is increased, this three-state distribution shifts toward the S_3_ state, with progressive disappearance of the S_2_ state. At ATP concentrations exceeding ∼ 2 µM, the distribution stabilizes in two states, a dominant S_3_ state plus a low level of the S_1_ state, which do not appear to respond further to increases in ATP concentration (Figure 4B). These results are in good qualitative agreement with previous smFRET measurements of SR1 in the presence of ATP (42).

**Figure 4.**
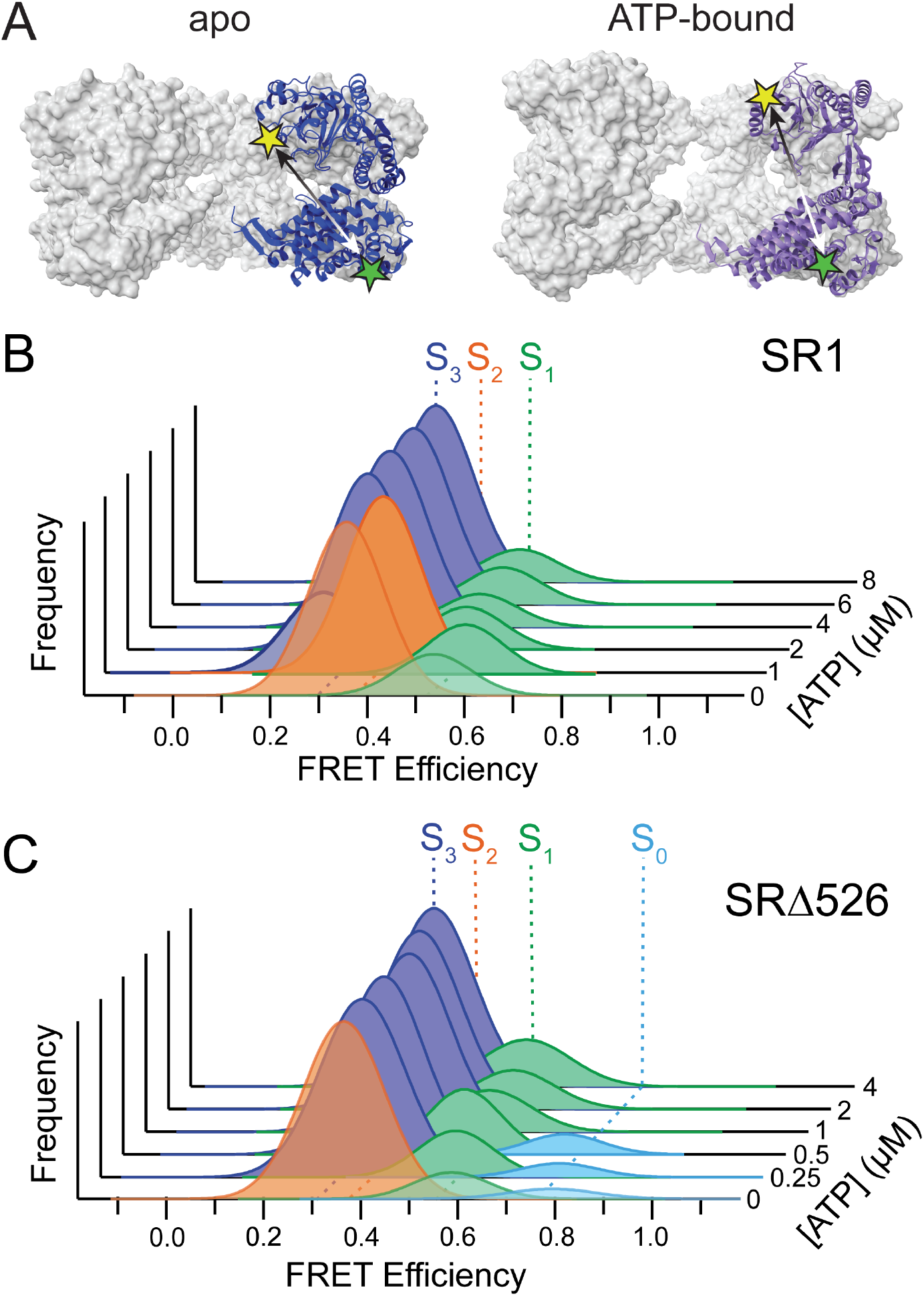
SRΔ526 populates more conformational states than SR1 in the absence of ATP. (A) Schematic of smFRET assay used to observe the conformational distribution of SR1 and SRΔ526 rings. Donor (Alexa488) and acceptor (Alexa594) probes were coupled to two engineered Cys residues (E255C and D428C) on one subunit of either an SR1 or SRΔ526 ring (42). Probe attachment positions are illustrated by colored stars on a single ring of either apo GroEL (PDBID: 1GR5) or ATP-bound GroEL (PDBID: 4AB3). The indicated shift in relative position (∼ 12 Å; double-headed arrows) sets the likely maximal distance change of the apical domains during this experiment. The smFRET efficiencies observed for SR1 (B) and SRΔ526 (C) at different ATP concentrations were fit to a minimal multi-state Gaussian distribution model. In the absence of ATP, SR1 is well described by two states (S_1_, S_2_), while SRΔ526 populates three (S_0_, S_1_, S_2_). Initial titration of SR1 with ATP (B) results in the appearance of a third, low FRET state (S_3_). At higher ATP concentrations, the S_2_ state disappears and the distribution becomes dominated by the low FRET S_3_ state. Addition of ATP to SRΔ526 results the apparent disappearance of the S_2_ state and its replacement by the S_3_ state at the lowest ATP concentrations examined, with the high-FRET S_0_ state persisting until the ATP concentration reaches ∼ 0.5 µM. At higher ATP concentrations the SRΔ526 micro-state distribution is very similar to that of ATP-saturated SR1. In all cases, samples were supplemented with an ATP regeneration system to maintain constant ATP concentrations during measurement.

Removal of the C-terminal tails alters the basal micro-state distribution of the ring and its response to ATP. When SRΔ526 was examined in the absence of ATP, three micro-states are observed (Figure 4C). Two of these states (S_1_ and S_2_) display FRET efficiencies similar to those observed with SR1, while the third, high-FRET state observed (S_0_) is not seen with SR1 (Figure 4C). The appearance of the S_0_ micro-state with SRΔ526 suggests this conformation is only accessible in the absence of the C-terminal tails. Very low concentrations of ATP (250-500 nM) result in the apparent total disappearance of the S_2_ state and its replacement by the S_3_ state. At the same time, the level of the S_2_ state increases modestly while the level of the high-FRET S_0_ state remains unchanged. Once the ATP concentration exceeds 0.5 µM, the S_0_ state disappears and the SRΔ526 distribution becomes essentially identical to that of SR1, with a dominant S_3_ state and lower level of the S_1_ state.

To gain greater insight into the underlying dynamics of these micro-state distributions, we applied photon-by-photon hidden Markov Modeling (H2MM) (67, 68). In H2MM, fluctuations in the observed FRET efficiency during each single molecule burst (in excess of measurement noise) are presumed to originate from Markovian transitions between distinct states (67, 68). H2MM thus provides a second method to characterize the minimal number of states needed to describe a data set, while also permitting measurement of the transition times between resolvable micro-states. In the absence of ATP, H2MM analysis of SR identifies two micro-states, consistent with Gaussian peak fitting of the FRET efficiency histograms (Figure 5A and Figure S3). Additionally, both burst variance analysis (BVA; Figure S3A) and dwell-based H2MM modeling (Figure 5A and Figure S3F) suggest modest interconversion dynamics between these two micro-states that occur on the 2-6 msec time scale.

**Figure 5.**
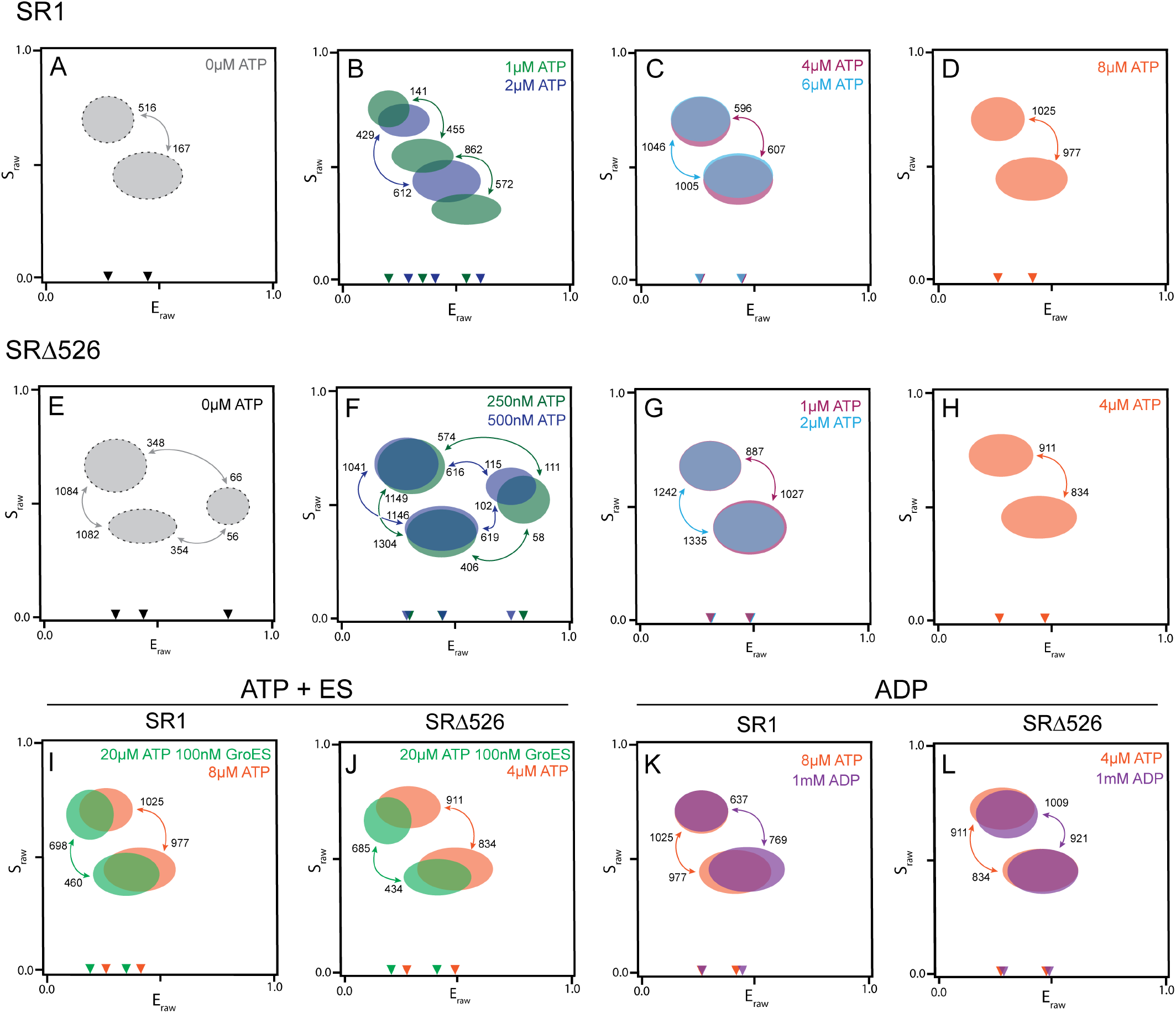
Deletion of the C-terminal tails increases micro-state dynamics of the SRΔ526 ring. Time-dependent fluctuations in smFRET efficiency for SR1 (A-C) and SRΔ526 (D-G) were modeled as Markovian state transitions using photon-by-photon H2MM analysis (mpH2MM) (67). The simplified distributions shown illustrate the approximate 0.95 population boundaries of each H2MM state identified (see Supplemental Information). Arrows indicate transitions between states that are observable by H2MM and the numbers next to each arrowhead specify the observed transition rate constants (s^-1^) for formation of the state to which the arrows point along the given path. (A) For apo SR1 (0 µM ATP), H2MM identifies two well-populated micro-states that interconvert on the 2-6 msec time scale. (B) Low concentrations of ATP (1 µM) result in an expansion of the number of SR1 micro-states (from two to three) and a decrease in the interconversion times for some transitions. (C) Higher ATP concentrations (2-8 µM) reduce the number of SR1 micro-states back to two and result in a progressive decrease in interconversion times from 1.5-2 msec to 0.9-1 msec. (D) SRΔ526 populates three distinct micro-states in the absence of ATP, including a high-FRET state not observed with SR1, which display a broad range of interconversion times from 16-17 msec to 0.9 msec. (E-F) Addition of low ATP concentrations (250-500 nM) results in a decrease in the interconversion time for several micro-states for SRΔ526 without an observable change in the number of detectable states. (G-H) Higher concentrations of ATP result in a two state distribution for SRΔ526. (I-J) Addition of GroES and ATP yield a two-state distribution for both SR1 and SRΔ526 that are similar to one another. (K-L) Addition of saturating concentrations of ADP (1 mM) to both SR1 and SRΔ526 yields two-state distributions that are slightly shifted to higher mean efficiency values compared to those seen at elevated ATP concentrations (C-D and G-H).

Addition of low ATP concentrations (1 µM) results in an expansion of the number of SR1 micro-states, along with the appearance of faster interconversion dynamics (Figure 5B). While some transitions appear to remain on the 2-6 msec time scale, others approach 1 msec. These observations are consistent with Gaussian peak fitting of the smFRET efficiency histogram at 1 µM ATP (Figure 4B), which identified three micro-states for the SR1 ring. As the ATP concentration is increased further, the number of SR1 micro-states shrinks, collapsing back to only two states by 2 µM ATP and remaining there as the ATP concentration is increased further (Figure 5B-D and Figure S4). Interestingly, while the distribution of states at 2 µM is very similar to that observed at higher ATP concentrations, transitions between these states become more rapid. Increasing the ATP concentration from 2 µM to 4 µM reduces the micro-state interconversion time from ∼ 2 msec to 0.9-1 msec (Figure 5B-D).

Application of H2MM to apo SRΔ526 demonstrates that C-terminal tail removal permits the ring to intrinsically populate a greater number of more rapidly interconverting micro-states than apo SR1 (Figure 5E and Figure S5). Three micro-states are resolved for SRΔ526 by H2MM in the absence of ATP, including the unexpected high-FRET state seen in Gaussian peak fitting of the smFRET efficiency histograms (Figure 4C). While the FRET efficiencies of the other two micro-states are similar to those observed with SR1, their positions on stoichiometry (S) vs efficiency (E) plots are shifted slightly (Figure 5A versus Figure 5E). The pattern of interconversion dynamics for the apo SRΔ526 distribution is also notably distinct from that of SR1 (Figure 5A). While the two apo SR1 micro-states interconvert on the 2-6 msec time scale, the similar SRΔ526 micro-states interconvert on the sub-msec time scale (∼ 0.9 msec). Interestingly, population of the high-FRET SRΔ526 micro-state occurs much more slowly (∼ 16 msec).

Addition of very low concentrations of ATP (0.25 - 0.5 µM) to SRΔ526 results in only small changes to the H2MM micro-state distribution (Figure 5E-F and Figure S6). However, while the fastest interconversion transitions remain on the sub-msec time scale, several SRΔ526 transitions that were slow in the apo state become more rapid upon ATP addition. The high-FRET SRΔ526 micro-state is also more highly populated at low ATP concentrations and displays faster formation and decay transitions (Figure 5E-F). As the ATP concentration increases to 1 µM, the SRΔ526 distribution collapses to just two micro-states with the disappearance of the high-FRET micro-state (Figure 5G). At all ATP concentrations examined above 1 µM, the observed E and S values of the remaining micro-states, as well as their interconversion dynamics, display values that are similar to those observed with SR1 at higher ATP concentrations (Figure 5G-H versus Figure 5C-D).

### Unexpected micro-state heterogeneity of the SR-ES complex is not impacted by the C-terminal tails

To determine whether the C-terminal tails influence the conformational dynamics of the GroEL ring upon GroES binding, we repeated our smFRET analysis in the presence of excess ATP and GroES. Prior observations have shown that SR1 binds ATP and GroES, executes a single hydrolytic turnover (t_1/2_ of approximately 8-10 sec at 25 °C), and stalls in a homogenous population of stable SR1-ADP-GroES complexes, which have a spontaneous disassembly half time of approximately 300 min under these conditions (19). Remarkably, H2MM analysis of SR1 in the presence of excess GroES and ATP resolves two smFRET micro-states, which appear to interconvert on the 1.5-2 msec time scale (Figure 5I). One of these states displays the lowest mean transfer efficiency of any state characterized in this study and possesses a more narrow spread of transfer efficiencies (Figure 5I). These features are consistent with a stable SR1-ADP-GroES complex in which the apical domains are bound and restrained by the associated GroES heptamer in a more open conformation than can be populated with saturating ATP alone (18, 43, 69). When SRΔ526 is mixed with excess ATP and GroES, it populates an essentially identical two-state distribution (Figure 5J), indicating that the C-terminal tails have no detectible impact on the conformational properties of the GroES-bound ring. However, the persistence of a second, higher-FRET micro-state, for both SR1 and SRΔ526, was unexpected.

To examine whether this surprising micro-state heterogeneity results from unanticipated GroES dissociation under these conditions, we examined both SR1 and SRΔ526 in the presence of a large excess of ADP (Figure 5K-L). More rapid dissociation of GroES could, in principle, yield sub-populations of ADP-bound rings at steady state that display distinct smFRET distributions. The micro-state distributions of SR1 and SRΔ526 in the presence saturating ADP (1 mM) are essentially identical (Figure 5K-L). These distributions are also composed of two micro-states with interconversion dynamics on the 1-1.5 msec time scale. The ADP-bound distributions for SR1 and SRΔ526 are slightly shifted to higher mean FRET efficiencies and, at least for SR1, possess slower interconversion dynamics, than ATP alone (Figure 5K). These observations are consistent with prior studies showing that ADP- and ATP-saturated GroEL rings are not conformationally equivalent, with the ADP-bound state likely more closed, on average (42, 43, 45). Importantly, these ADP-induced distributions are centered on mean FRET values that are distinct from those observed in the presence of GroES and ATP (Figure 5I-J). This strongly suggests that the unexpected micro-state heterogeneity and dynamics observed in the presence of GroES are not a consequence of unexpected dissociation of GroES from either SR1 or SRΔ526.

## DISCUSSION

In this study we examined how the GroEL C-terminal tails impact the allosteric response of a GroEL ring as it binds ATP. Using a pair of single ring variants (SR1 and SRΔ526) we discovered that C-terminal tail deletion results in a large (nearly 10-fold) increase in macroscopic ATP binding affinity for each subunit, along with a substantial increase in ATP binding cooperativity. Extraction of the per site ATP binding enthalpy (ΔH°) and entropy (ΔS°) using vT-nMS demonstrated that ATP binding to SR1 is almost completely entropy dominated until the ring is nearly filled (∼ SR1(ATP)_6_). Strikingly, removal of the C-terminal tails with SRΔ526 results in an altered thermodynamic signature for ATP binding, with a reduced entropic contribution to the first 3-4 ATPs and a near total inversion of energetic partitioning for the final thee binding steps, with the enthalpy change becoming dominant. Using smFRET, we found that the conformational distributions the GroEL apical domains, and their response to ATP, are also distinct in the presence and absence of the tails. While SR1 populates two slowly interconverting micro-states in the absence of ATP, the tailless SRΔ526 ring populates three states, two of which interconvert much more rapidly than is observed with SR1. Addition of ATP to both SR1 and SRΔ526 stimulates the rate of micro-state interconversion for both oligomers. However SR1, with its intact C-terminal tails, requires a higher concentration of ATP to achieve the same level of rapid dynamics as observed with SRΔ526.

In total, these observations suggest that the C-terminal tails play a central role in the allosteric design of the GroEL ring. We propose that the dynamics of the tails and their associated energy distributions are important for tuning the progression and sensitivity of the GroEL functional cycle (Figure 6). In the apo (T-like) macro-state, we hypothesize that the tails make direct, physical contact with the substrate-binding surfaces of the apical domains (helices H, I and underlying loop region including Y199 and Y203; (70)). This suggestion is consistent with the amphipathic character of the tails, the hydrophobic nature of the substrate protein-capturing apical domain surfaces and the length of the tails. Additionally, a prior molecular dynamics study suggested that, in the absence of nucleotides, interactions between the apical domains and C-termini are likely (71).

**Figure 6.**
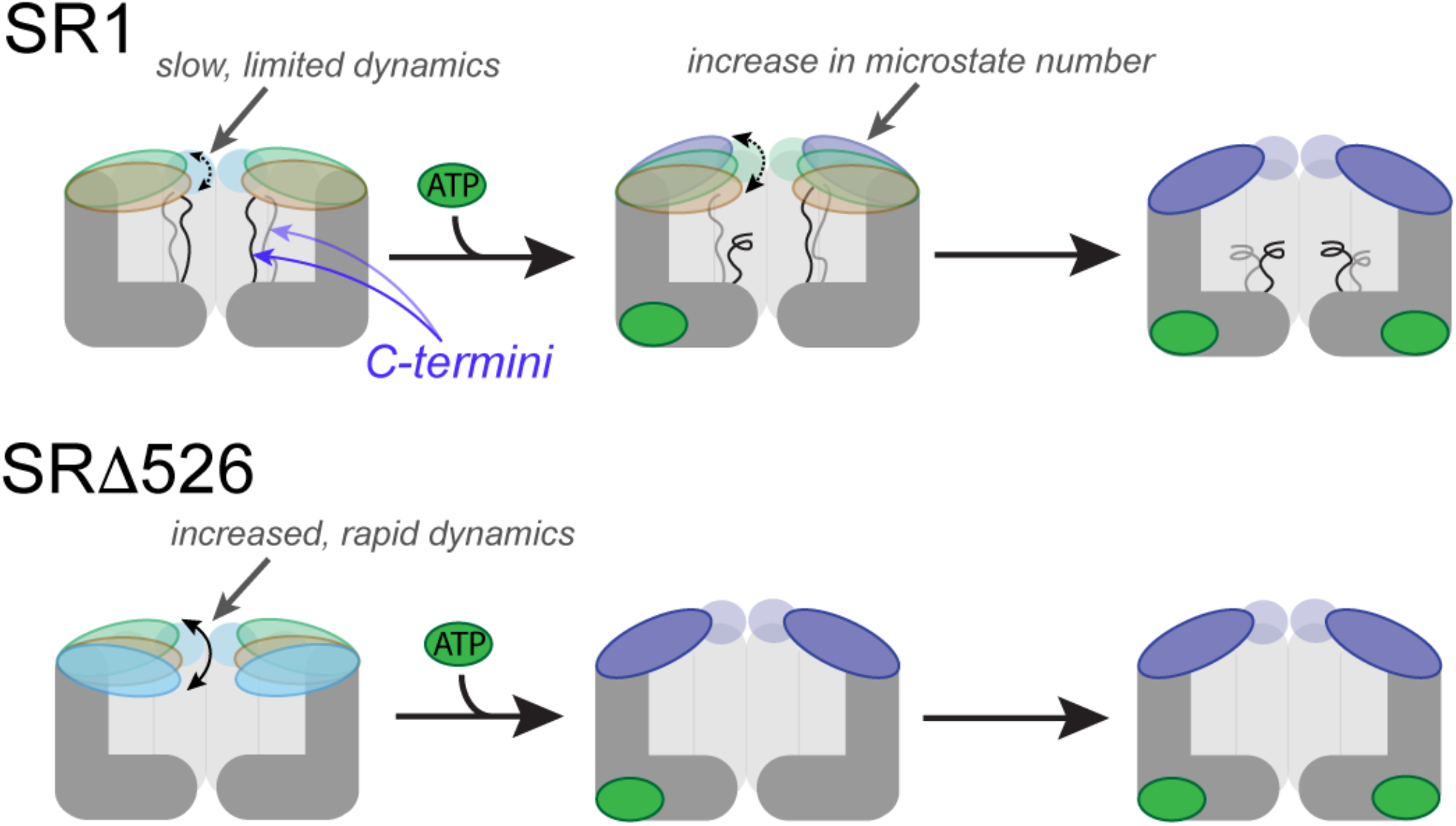
Model for the role of the intrinsically disordered C-terminal tails in regulating the allosteric progression of a GroEL ring. Schematic of an SR1 ring with intact C-termini is shown (top). The amphipathic and disordered C-terminal tails physically interact with the substrate binding surfaces of the SR1 apical domain, restricting both the micro-states that the apical domains can visit in the apo, T-like macro-state and how rapidly they can interconvert (dashed, double headed arrow). Because of the distance between the bottom of the GroEL cavity and the nominal position of the apical domains, this interaction requires either stretching or conformational restriction of the intrinsically disordered tails. Initial ATP binding to SR1 (above, center) initiates progressive release of the tails, increasing the number of micro-states available to the apical domains, the rate and which they can interconvert and the conformational entropy of both the tails and apical domains. The transition of the SR1 ring to its R-like macro-state occurs as the micro-state distribution is biased towards more open states as ATP fills the ring, eventually resulting in full release of the tails. In the absence of the C-terminal tails (SRΔ526, bottom), the apical domains of the apo, T-like macro-state intrinsically populate several micro-states that possess more rapid interconversion kinetics. C-terminal tail removal also appears to permit the SRΔ526 apical domains to swing through a larger total change in position, from a more closed state to a more open state. Without the bias provided by the C-terminal tails, the transition of the SRΔ526 ring to an R-like micro-state distribution begins to occur at the earliest stages of ATP binding.

We propose that interactions between the tails and apical domains adjust the ATP-binding cooperativity of the GroEL ring, inhibiting it from prematurely populating a GroES-binding state until a substrate protein has been captured (Figure 6). Mutual suppression of conformational dynamics in the tails and the apical domains is a key element of the model. For the tails to stretch and reach the apical surfaces located at the top of the GroEL ring, their conformational degrees of freedom must be reduced. At the same time, the load imposed by the tails on the apical domains limits the dynamics allowed around the apical-to-intermediate domain hinge point. The large-scale, rigid-body reorganization of the GroEL subunits that is triggered by ATP binding favors a more open conformational distribution of the apical domains (18, 43, 69, 72). These R-like micro-states inherently position the hydrophobic apical binding surfaces beyond the reach of the C-terminal tails, resulting in progressive tail release and a concomitant return of their intrinsic, dynamic disorder. At the same time, movement around the apical-to-intermediate hinge becomes less constrained as the tails dissociate, resulting in a more dynamic sampling of apical domain micro-state positions. These mutually limiting interactions between the apical domains and C-terminal tails thus encode an “entropic latch”, which is released as ATP binds. Importantly, without this entropic tuning, the GroEL ring appears to snap into a R-like micro-state distribution (e.g. SRΔ526) too easily. This model is also consistent with the role proposed for flexible or disordered domains in other allosteric systems (40, 73, 74).

While the conformational dynamics of the tails and apical domains play a pivotal role, changes in surface hydration are also likely important. The positive ΔC_p_ observed for most ATP binding steps during vT-nMS experiments (Table S1) is consistent with a nucleotide-induced increase in hydrophobic surface exposure. Interestingly, the progression of increased heat capacity, as a function of ring saturation, is more gradual for SR1 than SRΔ526 (Table S1). This observation is consistent with the reduced ATP binding cooperativity seen with SR1 (Figure 1). It is also consistent with the idea of a progressive disruption of tail-apical domain interactions, where hydrophobic apical surfaces that are freed of their interaction with the tails become more fully exposed to solvent as the ring opens.

Ring opening is ultimately required to form a closed GroEL-GroES complex, in which folding intermediates are encapsulated and folding initiated (19–21, 75). Prior studies have shown that this complex is highly structured and stable (18, 19, 76). Observation of apical domain dynamics in the SR1-GroES and SRΔ526-GroES complexes are, consequently, surprising and suggest an unexpected degree of micro-state heterogeneity. A previous smFRET study of SR1 observed similar dynamics in the SR1-GroES complex, which were attributed to periodic detachment and re-binding of a subset of apical domains to their GroES mobile loops (42). We believe that an alternative hypothesis is more likely. Released from the constraints of the second GroEL ring, the equatorial plate of the SR1 oligomer may populate structurally variable, and dynamic, conformational states not normally available in tetradecamer GroEL. If correct, the allosteric signal that is transmitted from an ATP-saturated ring through the equatorial ring-ring interface, which encodes the well-characterized negative ATP binding cooperativity between the rings (26–29), encounters no resistance in SR1 or SRΔ526. As a result, the equatorial domains of these single ring oligomers, which carry one of the FRET probes, may be capable of oscillating away from the GroES-bound apical domains, thereby contributing to the observed smFRET micro-state distributions. Indeed, the lowest FRET sub-state of the ATP-saturated SR1 and SRΔ526 rings, even in the absence of GroES, might already contain elements of this behavior (Figure 5). Consistent with this supposition, prior cryoEM studies demonstrated that an SR1-GroES complex can encapsulate much larger substrate proteins than tetradecamer GroEL, apparently as a direct consequence of increased structural plasticity of the SR1 equatorial domain contacts (77). Interestingly, the absence of the C-terminal tails (i.e. SRΔ526) appears to have no impact on this behavior, despite the attachment of the tails to the equatorial domains on the cavity interior (Figure 5).

Importantly, the C-terminal tails of GroEL are also involved in substrate protein capture and folding. We and others have shown that the amphipathic tails physically interact with non-native folding intermediates, both in an open, apo ring and within the GroEL-GroES complex (49, 51, 55, 56). We and others have also shown that removal of the tails reduces assisted folding efficiency, compromises substrate protein retention and suppresses an ATP-driven forced unfolding event that accompanies GroES encapsulation of substrate proteins (48–54). Combined with the results presented here, these observations suggest a unified model of C-terminal tail function, in which these intrinsically disordered domains act as a central allosteric integrator for the GroEL functional cycle. By binding to the apical domains of the apo GroEL ring, the tails reduce the chance that an empty ring, containing no substrate protein, can transition to the GroES acceptor state. While an empty complex will ultimately disassemble and reset during the next round of the cycle, excessive “misfires” of this type will inevitably reduce proteostatic efficiency and lead to compromised overall fitness. Additionally, stringent non-native substrate proteins are likely to display hydrophobic surfaces that interact much more strongly with the apical domains than do the tails. Folding intermediate binding would thus displace the tails from the apical domains and signal successful substrate protein capture. In turn, this would trigger a shift of the ring to a more highly cooperative ATP binding regime, promoting effective and near switch-like encapsulation of the substrate protein beneath GroES. In total, this sequence of events enforces an ordered and efficient binding of substrate protein, ATP and GroES and ensures a minimal likelihood of empty complex formation and wasted metabolic energy.

The entropic tuning model we propose is also consistent with recent re-formulations of allosteric theory, in which shifts in micro-state dynamics are proposed to be a unifying foundation of molecular allostery (34–37). In this view, the probabilistic events that underpin allosteric signaling can occur across times scales that range from nsec to sec, and can be based in either very fast amino acid side chain restructuring, moderately fast backbone re-organization and folding or classical rigid body re-orientation of well folded domains. In the model we propose here, fast conformational dynamics of the intrinsically disordered C-terminal tails, and their associated energetics, are utilized to control the slower and classically describable allosteric response (e.g. the T to R macro-state transition) of a GroEL ring. One of the challenges facing the modern dynamic view of allostery is understanding how conformational dynamics in complex oligomeric systems are coupled across functionally relevant time and length scales. We suggest that the C-terminal tails of GroEL provide one evolutionary solution to this problem. Given the frequency with which intrinsically disordered domains and tails seem to appear in other allosteric systems, particularly several of the other molecular chaperone families, a fascinating question is whether the conserved and disordered domains in these other systems play similar entropic turning roles.

## METHODS

### Protein Expression and Purification

SR1, SRΔ526, and the double Cys mutants of SR1(E225C/D428C) and SRΔ526(E225C/D428C) were purified as described previously (49, 50). Briefly, each protein was expressed from IPTG-driven promoters using *E. coli* BL21. Cells were pelleted by centrifugation, resuspended in lysis buffer (50 mM Tris pH 8.0, 0.5 mM EDTA, 2 mM DTT, 20% sucrose), shear ruptured using a Microfluidizer LM-20 (Microfluidics) and centrifuged (35,000x g) to removed cell debris. The clarified lysate was then loaded onto an Fast Flow Q (Cytiva) anion exchange column in 50 mM Tris pH 7.4, 0.5 mM EDTA, 2 mM DTT, which was developed using a linear NaCl gradient. SR1-or SRΔ526-containing fractions were combined and dialyzed against 50 mM Bis-Tris pH 6.0, 12.5% MeOH, 0.5 mM EDTA, 50 mM KCl, 2 mM DTT, then loaded onto a second, dedicated Fast Flow Q stripping column in 50 mM Bis-Tris pH 6.0, 12.5% MeOH, 0.5 mM EDTA, 2 mM DTT), which was developed using a linear NaCl gradient. SR1- and SRΔ526-containing fractions were combined and further stripped of adhering contaminants by batch washing (with gentle agitation under an argon atmosphere) with Affigel Blue resin (50-100 mesh, 150-300µm; BioRad) for 15-20 hours at 4 °C in 50 mM Bis-Tris pH 6.0, 12.5% MeOH, 0.5 mM EDTA, 2 mM DTT. Single ring oligomers were then separated from contaminating endogenous GroEL tetradecamer using an S300 gel filtration column equilibrated in 25 mM Tris pH 7.4, 100 mM KCl, 0.5 mM EDTA, 2 mM DTT). Samples of purified SR1 and SRΔ526 were concentrated to 10-20 mg/ml, supplemented with glycerol to a final concentration of 15% (v/v), snap frozen in liquid N_2_ and stored at -80 °C.

### Native Ion Mass Spectrometry

All reagents used for MS analysis, including ethylenediaminediacetic acid (EDDA), ATP, and magnesium acetate (MgAc) were purchased from Sigma-Aldrich (St. Louis, MO) and were dissolved in LC-MS grade deionized water. Protein samples were diluted into 200 mM EDDA, pH 6.3, 1 mM MgAc for all MS experiments and ATP was prepared from anhydrous powder immediately prior to use. Stock samples of either SR1 and SRΔ526 were quick thawed and buffer exchanged into MS EDDA buffer immediately prior to use using MicroBio P-6 gel spin columns.

The sample temperature was controlled during variable temperature native ion mass spectrometry (vT-nMS) experiments using a custom-built variable temperature device incorporated into the nano-ESI emitter, as described previously (59, 78). This device possesses a calibrated temperature error of +/- 2 °C. The temperature range employed during these experiments was limited to 4-35°C. Working ATP solutions were prepared by diluting concentrated nucleotide stocks into the same MS EDDA buffer at twice the final target nucleotide concentration. SR1 or SRΔ526 samples were then mixed with ATP using a 1:1 mixing protocol to achieve the indicated protein:ATP ratios prior to sample loading into the temperature-controlled MS emitter. All samples were incubated at each temperature for 1 minute prior to injection into an Thermo Q Exactive UHMR (ultra-high mass range) hybrid quadrupole Orbitrap mass spectrometer. For all experiments, the scan resolution was set to 12500 using a single micro-scan with 100 scans collected for each spectrum. The MS inlet capillary temperature set to 120°C with in source trapping set to -250V and the HCD energy was set to 250.

Processing of raw MS data for species mass and abundance was carried out using UniDec (79). The observed abundances of each SR1 and SRΔ526 ligation state, at each ATP concentration and temperature, were fit to a sequential binding model to extract observed macroscopic association constants for each ligation state (65). The temperature dependence of each binding constant was then fit to a non-linear variant of the van’t Hoff equation to extract enthalpic (ΔH), entropic (-TΔS) and heat capacity (ΔC_p_) changes associated with the formation of each ligation state (65).

### Protein Labeling

SR1(E225C/D428C) and SRΔ526(E225C/D428C) were fluorescently labeled essentially as previously described with some modifications (42). Briefly, samples of SR1(E225C/ D428C) or SRΔ526(E225C/D428C) were buffer exchanged into reaction buffer (50 mM Tris pH 7.4, 100 mM KCl, 0.5 mM EDTA, 0.1 mM TCEP) using a PD-10 desalting column. Solutions of AlexaFluor-488-maleimide (AF488) and AlexaFluor-594-maleimide (AF594; ThermoFisher) were prepared from dry powder in anhydrous N,N-dimethylformamide immediately prior to use. Protein samples (5-10 mg/ml final protein concentration) were first mixed with (100-200 µM final dye) AF594 (final AF594:site mixing ratio of 1.2:1) for 45 min at 25 °C with constant stirring in the dark. Samples were then supplemented with AF488 (100-200 µM final dye) AF488 mixing ratio of 1.2:1) and incubated in the dark at 25 °C for an additional 45 min with stirring. The reaction was quenched by the addition of 5 mM glutathione followed by an additional incubation in the dark for 30 min. Unreacted dye was removed using a PD-10 desalting column equilibrated with storage buffer (50 mM Tris pH 7.4, 100 mM KCl, 0.5 mM EDTA, 2 mM DTT). Concentrated samples of labeled SR1(E225C/D428C) or SRΔ526(E225C/ D428C) were then mixed into disassembly buffer (4 M deionized urea, 50 mM Bis-Tris pH 6.0, 1 mM DTT) and incubated at 25 °C for 30 minutes to dissociate SR1 and SRΔ526 into monomers. Dissociated samples were loaded onto a MonoQ 5/50 column (Cytiva) equilibrated in the same buffer and developed using a linear NaCl gradient. Purified double-labeled SR1(E225C/D428C) or SRΔ526(E225C/D428C) monomers were collected and immediately mixed with excess (1:100) unlabeled and dissociated SR1 or SRΔ526 monomers, which had been subjected to the same urea buffer dissociation protocol above. Re-assembly of SR1 or SRΔ526 rings containing, at most, a single, labelled SR1(E225C/D428C) or SRΔ526(E225C/D428C) subunit was carried out by overnight dialysis of mixed samples into reassembly buffer (50 mM Tris pH 7.4, 0.6 M NH_4_SO_4_, 10 mM MgCl_2_, 2-5 mM ADP, 5 mM DTT) in the dark at 23 °C. Reassembled SR1 and SRΔ526 samples were centrifuged at 100,000x g to remove residual aggregates that were observed to form during reassembly. Final samples were concentrated to 10-20 mg/ml, supplemented with glycerol to a final concentration of 15% (v/v), snap frozen in liquid N_2_ and stored at -80 °C.

### Single Molecule Förster Resonance Energy Transfer

smFRET experiments were carried out using extensively cleaned and BSA-passivated round glass No. 1.5 coverslips (Ted Pella) that were mounted and secured in a multi-well, custom-built PEEK coverslip holder. Once loaded, the coverslip holder was mounted to the stage of a four-channel, inverted single molecule Luminosa confocal microscope (PicoQuant). For each measurement, samples of labeled SR1 or SRΔ526 (containing no more than one double-labeled subunit) were diluted to a final concentration of 50-100 pM rings in 25 mM Tris HCl pH 7.4, 10 mM Mg(OAc)_2_, 10 mM KOAc buffer supplemented with either 25 nM unlabeled SR1 or SRΔ526 oligomer to stabilize ring assembly. Sample droplets (∼ 10 µl) containing the indicated concentrations or SR1 or SRΔ526, mixed with ATP, ADP or ATP + GroES were placed on the top of the coverslip and the droplet was centered on a 60x/1.2 water immersion objective (Olympus UPlanSApo), with a typical focal position of 7 µm into the droplet (past the upper coverslip surface). All samples containing ATP were also supplemented with an ATP regeneration system (30 units/mL creatine kinase and 3 mM creatine phosphate) to maintain constant ATP levels during measurement and prevent the accumulation of ADP. A humidifier cap was placed over the sample droplet to prevent evaporation during acquisition. Fluorescence burst data was collected from freely diffusing SR1 and SRΔ526 oligomers using pulsed interleaved excitation (PIE) (80–82) at 23 °C, using a 485 nm donor excitation laser (25 µW) and a 560 nm acceptor excitation laser (35 µW), alternately pulsed at 20 MHz each. Time-correlated single photon counting (TCSPC) data was collected for the donor emission using a 520 ± 35 nm bandpass filter and acceptor emission was collected using a 561 long pass filter combined with a 595 ± 40 bandpass filter. Data was continuously collected for 45 min to 1 hr for each measurement and each condition was replicated a minimum of three, independent times. For all data records, the observed burst frequency was typically between 1-2 bursts/sec and the observed burst width (FWHM) was 5 ± 2 msec. Background (buffer) signal levels were typically 0-0.08 counts/sec.

Corrected smFRET efficiencies used for Gaussian multi-peak fitting were calculated from raw, 2-channel correlated fluorescence burst data using the PicoQuant Luminosa software. The lower burst detection threshold was set at 1000 counts/sec and bursts were amplitude filtered using a minimum threshold of 100 total counts per burst (donor plus acceptor). Accepted bursts typically ranged from 50-200 counts per channel. The smFRET correction factors, (1) **α** - for leakage of the donor emission into acceptor detection channel, (2) ***δ*** - for direct excitation of the acceptor dye by the donor excitation laser, (3) **β** - normalization of donor and acceptor excitation intensities and emission cross-sections and (4) ***γ*** - normalization of fluorescence quantum yields and microscope channel detection efficiencies, were internally calculated using the Luminosa data acquisition software from in-sample donor-only and acceptor-only populations (82). Donor and acceptor laser powers were adjusted prior to full acquisition to minimize the **β** and ***γ*** correction factors. Corrected, amplitude filtered bursts from a minimum of three replicates were combined to yield total event data sets of 6000-10000 bursts (∼ 4 hours of total data collection). Multi-peak Gaussian peak fitting of the observed smFRET efficiency histograms was performed using the data processing package IgorPro (Wavemetrics) and its Gaussian multi-peak function. Gaussian components were incrementally added to the fit model until the observed fit residuals and reduced **χ**^2^ fit were minimized.

### Hidden Markov Modeling of smFRET data

Identification, processing and photon-by-photon Hidden Markov Modeling (H2MM) of smFRET data was performed using a combination of the open source Python packages FRETBursts (https://github.com/harripd/FRETBursts) and H2MM_C (https://github.com/ harripd/H2MMpythonlib) (67). Raw, time-tagged photon data was converted to phHDF5 (https://photon-hdf5.github.io/) format prior to loading into FRETBursts (<u>https://</u> <u>github.com/OpenSMFS/FRETBursts</u>) (83, 84). Burst events were identified using an all channel burst search algorithm, with background count levels calculated for each channel using a 30 sec sliding window across each data record. Bursts were identified using a photon count rate of m = 10 consecutive photons and a discrimination level of F ≥ 6 times the local background count rate. Identified bursts were amplitude filtered using a minimum threshold of 50 total photons (from both donor and acceptor data streams) per burst. Filtered burst data sets were then imported into a Jupyter notebook (https://zenodo.org/records/5566886) configured to implement the H2MM_C processing package to conduct both single parameter H2MM (spH2MM) (68) and multi-parameter H2MM (mpH2MM) (67). Because prior studies indicated that mpH2MM yields a more robust characterization of the state distribution and interconversion kinetics (67), the H2MM models specified by mpH2MM were typically used for data interpretation. Model performance as a function of increasing complexity was evaluated using both the (C) integrated complete likelihood (ICL) and (D) Bayesian information criterion (BIC), with the best fit model expected to minimize these statistics.

## SUPPORTING INFORMATION

Supporting data figures (S1-S6) and table (S1).

## ACKNOWLEDGEMENTS

We would like to Dr. Chavela Carr for editorial contributions to the manuscript and members of the Rye and Russell laboratories for their valuable suggestions and discussions. We would also like to thank Dr. Paul Harris for very helpful discussions and guidance in the configuration and use of mpH2MM and FRETBursts.

## AUTHOR CONTRIBUTIONS

Conceptualization: MEP, DHR, HSR

Methodology: MEP, KAE, HMS, DHR, HSR

Investigation: MEP, KAE, HMS

Formal Analysis: MEP, KAE, HMS

Validation: MEP, KAE, HMS

Supervision: DHR, HSR

Funding Acquisition: DHR, HSR

Writing - Original Draft: MEP, HSR

Writing - Review and Editing: MEP, DHR, HSR

## DATA SHARING

All data are available in the main text or the supplemental methods. Raw data sets available upon reasonable request from the corresponding authors.

