## Supplemental Information for "Regulation of a Classical Allosteric Molecular Machine by an Intrinsically Disordered Domain: the C-termini of GroEL"

**Table S1.** Site-resolved change in heat capacity (ΔCp) for SR1 and SRΔ526 as a function of ligation state.

| ATPs bound | SR1 $\Delta C_p$ ($kJ \cdot mol^{-1} \cdot K^{-1}$) | SRΔ526 $\Delta C_p$ ($kJ \cdot mol^{-1} \cdot K^{-1}$) |
|---|---|---|
| 1 | 0.27 ±0.10 | 0.62 ± 0.65 |
| 2 | 0.17 ± 0.11 | 0.73 ± 0.27 |
| 3 | 0.27 ± 0.22 | 0.89 ± 1.14[a] |
| 4 | 0.44 ± 0.15 | 2.07 ± 2.82[a] |
| 5 | 0.66 ± 0.07 | 3.39 ± 1.58[a] |
| 6 | 0.55 ± 0.27 | 1.63 ± 0.66 |
| 7 | 0.31 ± 0.14 | -0.58 ± 0.25 |

[a] Due to the higher cooperatively displayed by SRΔ526, the population of intermediate ligation states n = 3-5 was too low at most ATP concentrations for robust fitting of $\Delta C_p$ for these binding steps.

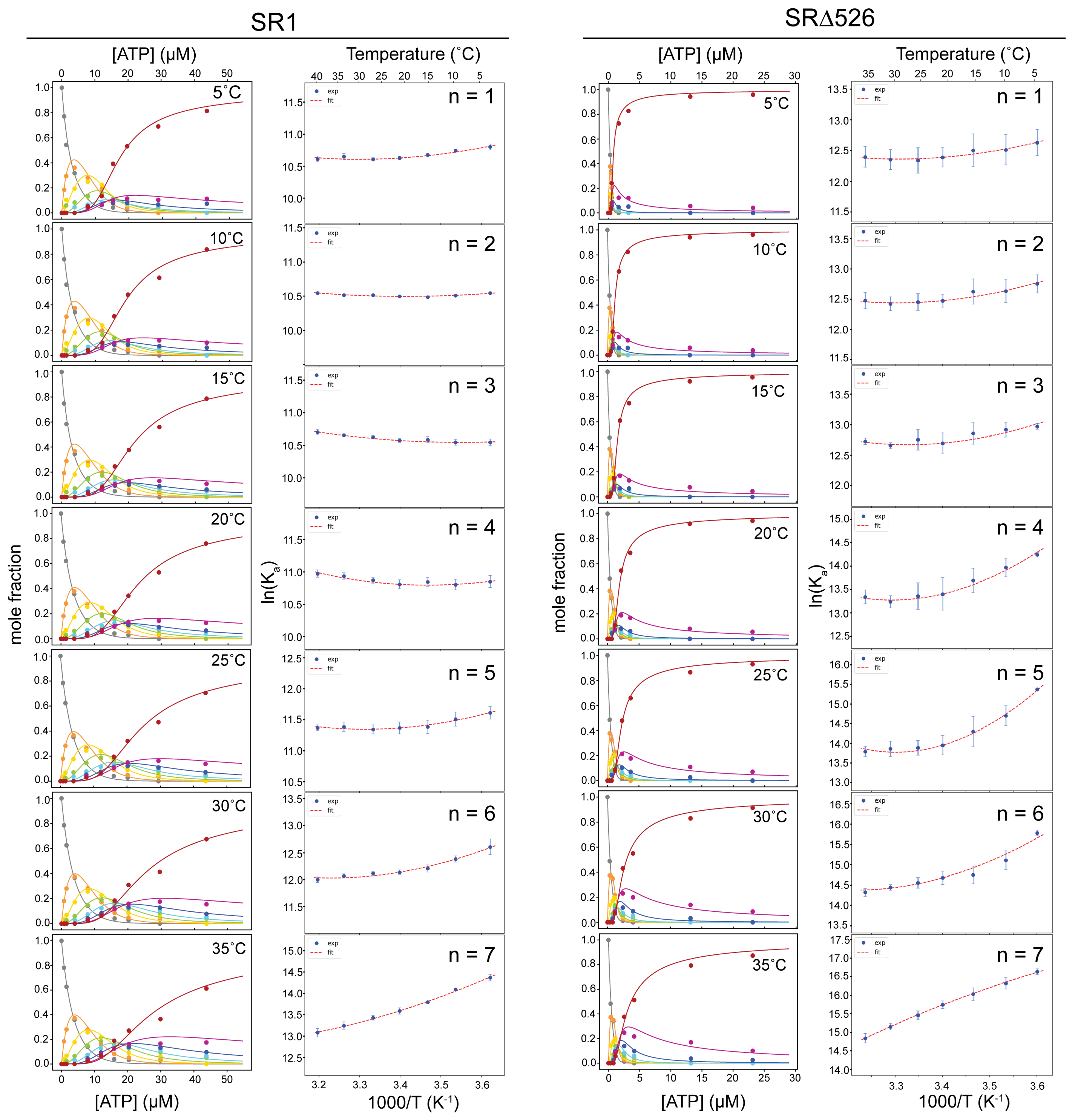


**Figure S1. Site-resolved binding of ATP to SR1 and SRΔ526 as a function of temperature.** Representative mole fraction plots for SR1-ATP$_n$ (first column) and SRΔ526-ATP$_n$ (third column) for n = 1-7 bound ATPs at different temperatures. Temperature-dependent mole fraction data was fit to a macroscopic sequential binding model (solid lines) to extract apparent association constants ($K_a$) associated with the formation of each ligation state (61, 79). Colors corresponds to the number of ATP bound each oligomer: n=0, grey; n=1, orange; n=2, yellow; n=3, green; n=4, light blue; n=5, dark blue; n=6, magenta; n=7, red. The temperature dependence of each ligation state association constant for SR1 (second column) and SRΔ526 (fourth column) was fit to a non-linear form of the van't Hoff equation (dotted lines) to determine the enthalpy (ΔH), entropy (ΔS) and heat capacity ($\Delta C_p$) changes associated with each ATP binding event (61, 79).

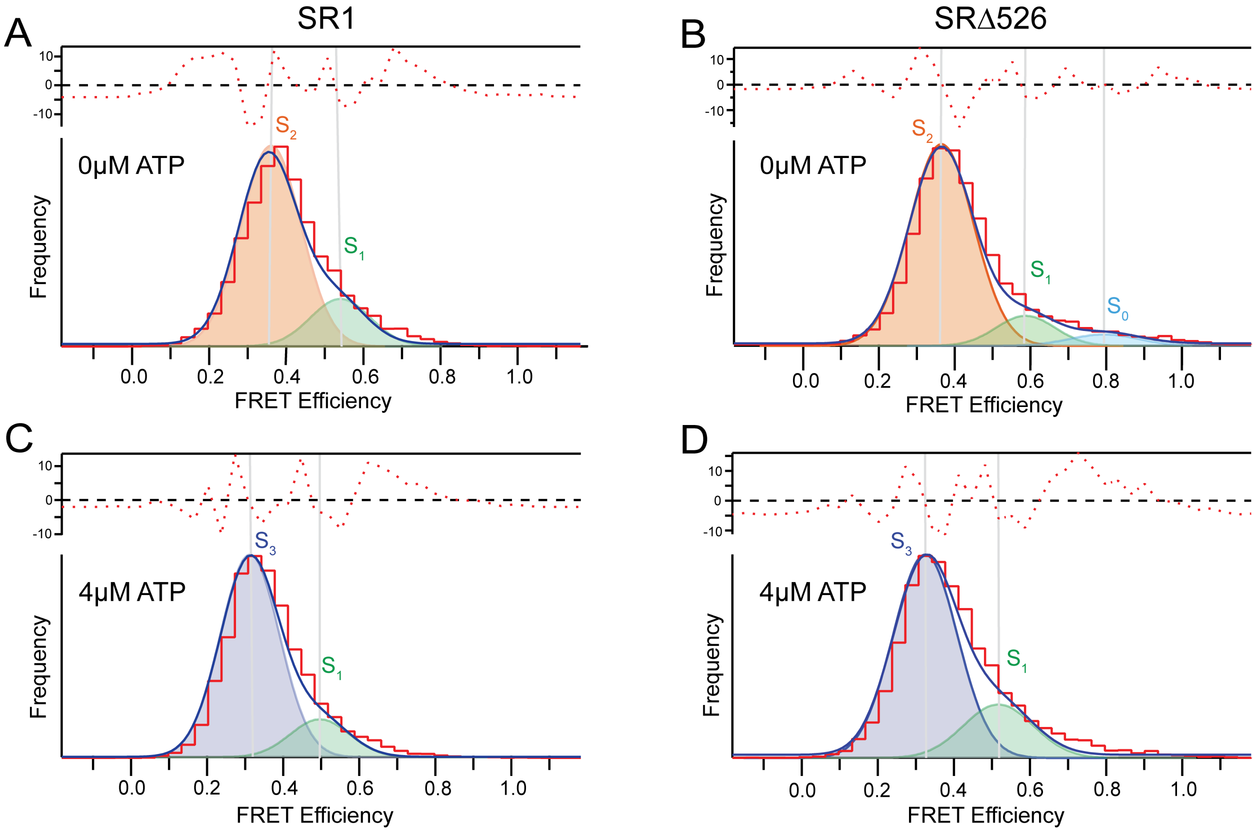


**Figure S2. Gaussian peak fitting of smFRET efficiency for SR1 and SRΔ526 identifies micro-states that change in response to ATP.**

Following burst identification, smFRET data for SR1 in the (A) absence and (C) presence of saturating ATP (4 μM) was binned (n = 39 bins) between efficiency (E) values of 0.0 to 1.0 (red histogram line) and fit to a minimal multi-component Gaussian model (black). For apo SR1, two components are required, with centroid E values of 0.37 ($S_2$, orange) and 0.52 ($S_1$, green). In the presence of 4 μM ATP, the SR1 data shifts to lower FRET efficiencies, requiring two fitting components with centroid E values of 0.31 ($S_3$, purple) and 0.50 ($S_1$). (B) In the absence of ATP, SRΔ526 FRET efficiency histograms require a three component fit, with centroid E values of 0.37 ($S_2$), 0.55 ($S_1$), and 0.8 ($S_0$). (D) Addition of 4 μM ATP to SRΔ526 results in a shift of the efficiency histogram to lower FRET efficiencies, similar to that of SR1, requiring two components with centroid E values of 0.31 ($S_3$) and 0.51 ($S_1$). Residual plots for each multi-component Gaussian fit are shown above each histogram (dotted lines).

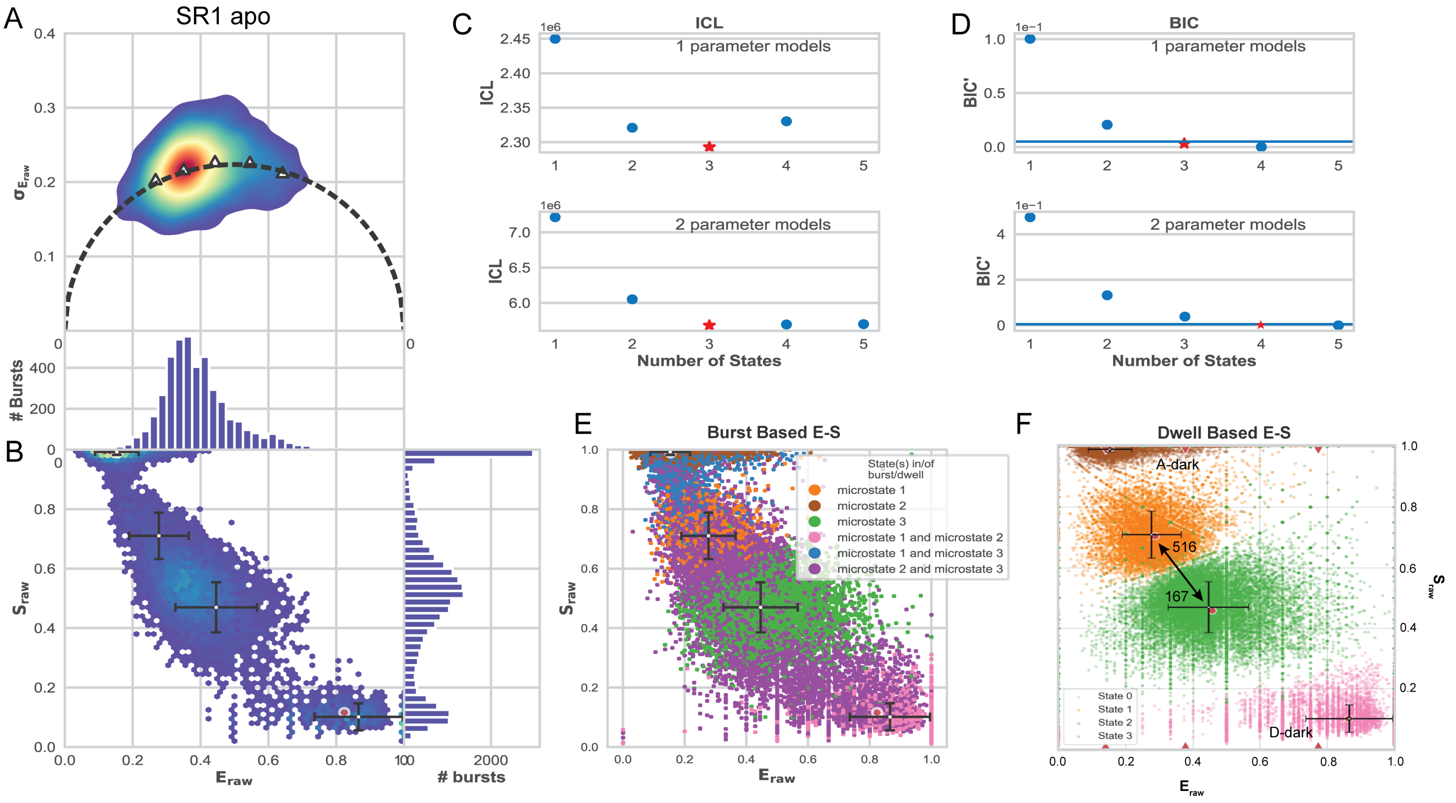


**Figure S3. Photon-by-photon H2MM analysis of apo SR1.**
(A) smFRET burst variance analysis (BVA) of 4337 bursts for apo SR1 identified by an all channel burst search (ACBS) (83, 84). (B) Stoichiometry (S) versus FRET efficiency (E) plot of raw smFRET burst data. H2MM analysis was performed using spH2MM (C and D, upper) and mpH2MM (C and D, lower) algorithms (67). Model performance as a function of increasing complexity was evaluated using (C) integrated complete likelihood (ICL) and (D) Bayesian information criterion (BIC) methods. The 'star' in each plot highlights the minimum number of states identified by each analysis. (E) Burst-based and (F) dwell-based plots of smFRET bursts colored by identified micro-state. The centroid of each micro-state distribution is shown (red and white dots), with error bars representing the standard deviation of the S and E values from Viterbi analysis. In the absence of ATP, SR1 is well described by two FRET micro-states, in addition to a donor-dark (D dark) and an acceptor-dark (A dark) state. Interconversion of the FRET micro-states, indicated by the double-headed arrow and observed rate constants ($s^{-1}$) in (F), occurs on the 2-6 msec time scale. For presentation clarity the interconversion pathways into and out of the D-dark and A-dark states are not shown.

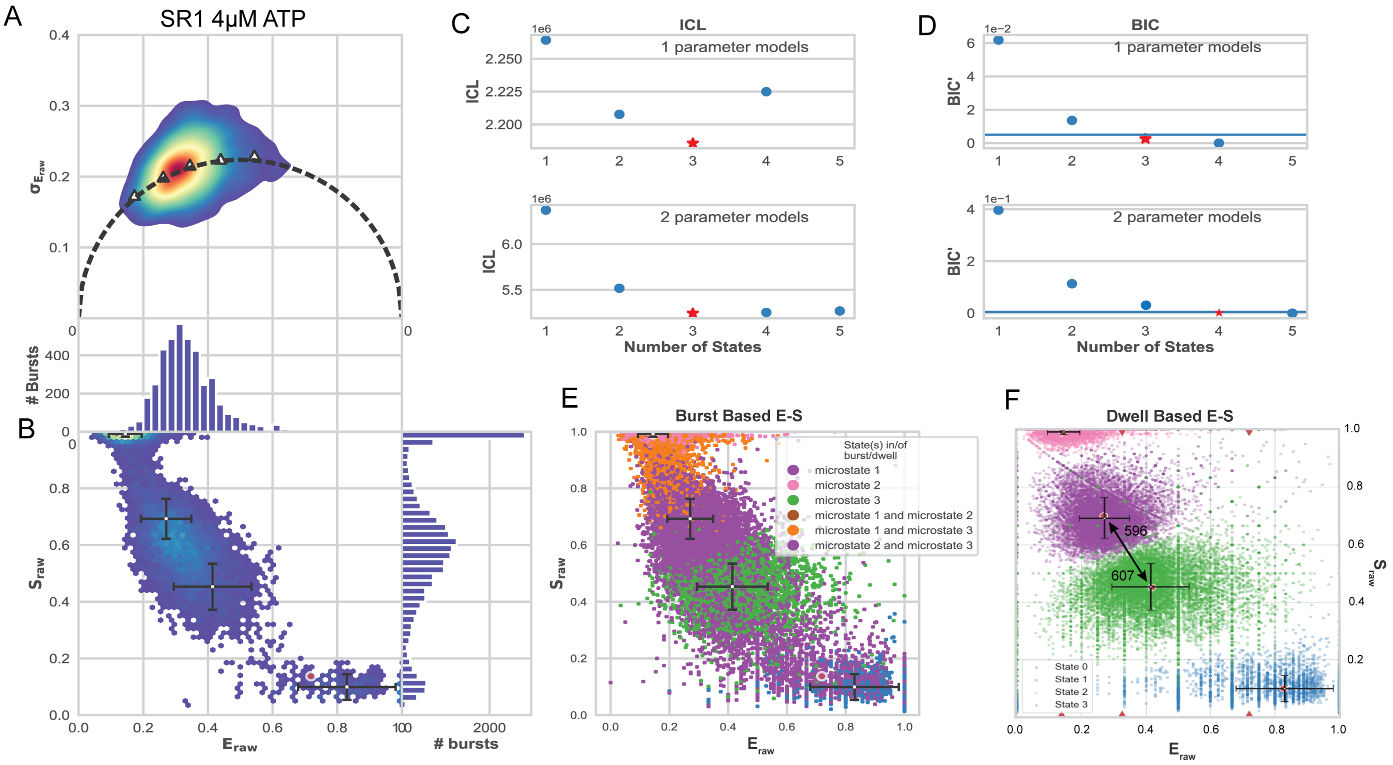


**Figure S4. Photon-by-photon H2MM analysis of SR1 plus 4 μM ATP.**

(A) smFRET burst variance analysis (BVA) of 3991 bursts for SR1 in the presence of 4 μM ATP identified by an all channel burst search (ACBS) (83, 84). (B) Stoichiometry (S) versus FRET efficiency (E) plot of raw smFRET burst data. H2MM analysis was performed using spH2MM (C and D, upper) and mpH2MM (C and D, lower) algorithms (67). Model performance as a function of increasing complexity was evaluated using (C) integrated complete likelihood (ICL) and (D) Bayesian information criterion (BIC) methods. The 'star' in each plot highlights the minimum number of states identified by each analysis. (E) Burst-based and (F) dwell-based plots of smFRET bursts colored by identified micro-state. The centroid of each micro-state distribution is shown (red and white dots), with error bars representing the standard deviation of the S and E values from Viterbi analysis. In the presence of 4 μM ATP, SR1 is again described by two FRET micro-states, in addition to a donor-dark (D dark) and an acceptor-dark (A dark) state. Interconversion of the FRET micro-states, indicated by the double-headed arrow and observed rate constants ($s^{-1}$) in (F), now occurs on the 1-2 msec time scale. For presentation clarity the interconversion pathways into and out of the D-dark and A-dark states are not shown.

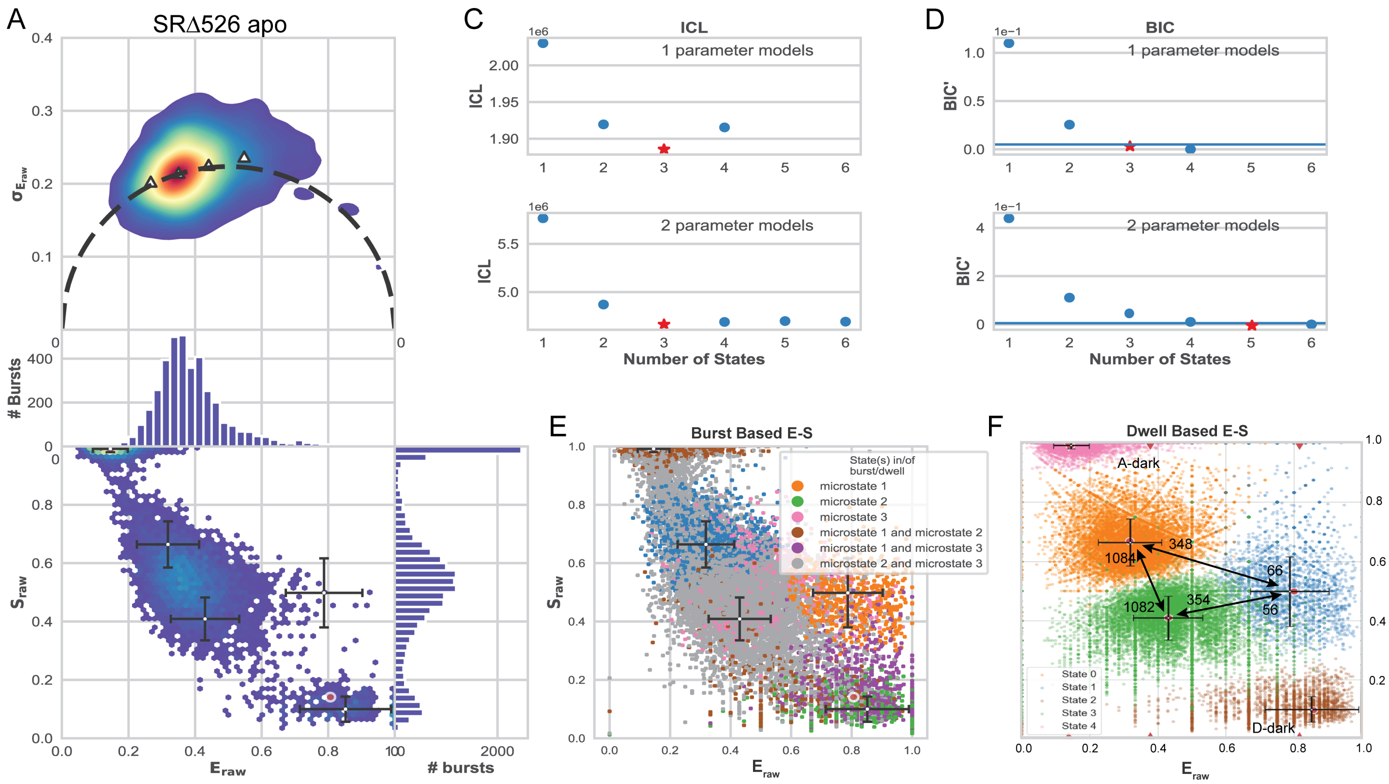


**Figure S5. Photon-by-photon H2MM analysis of apo SRΔ526.**

(A) smFRET burst variance analysis (BVA) of 3850 bursts for apo SRΔ526 identified by an all channel burst search (ACBS) (83, 84). (B) Stoichiometry (S) versus FRET efficiency (E) plot of raw smFRET burst data. H2MM analysis was performed using spH2MM (C and D, upper) and mpH2MM (C and D, lower) algorithms (67). Model performance as a function of increasing complexity was evaluated using (C) integrated complete likelihood (ICL) and (D) Bayesian information criterion (BIC) methods. The 'star' in each plot highlights the minimum number of states identified by each analysis. (E) Burst-based and (F) dwell-based plots of smFRET bursts colored by identified micro-state. The centroid of each micro-state distribution is shown (red and white dots), with error bars representing the standard deviation of the S and E values from Viterbi analysis. In the absence of ATP, SRΔ526 is well described by three FRET micro-states, in addition to a donor-dark (D dark) and an acceptor-dark (A dark) state. Interconversion of the FRET micro-states, indicated by the double-headed arrow and observed rate constants ($s^{-1}$) in (F), occurs on the 0.9-18 msec time scale. For presentation clarity the interconversion pathways into and out of the D-dark and A-dark states are not shown.

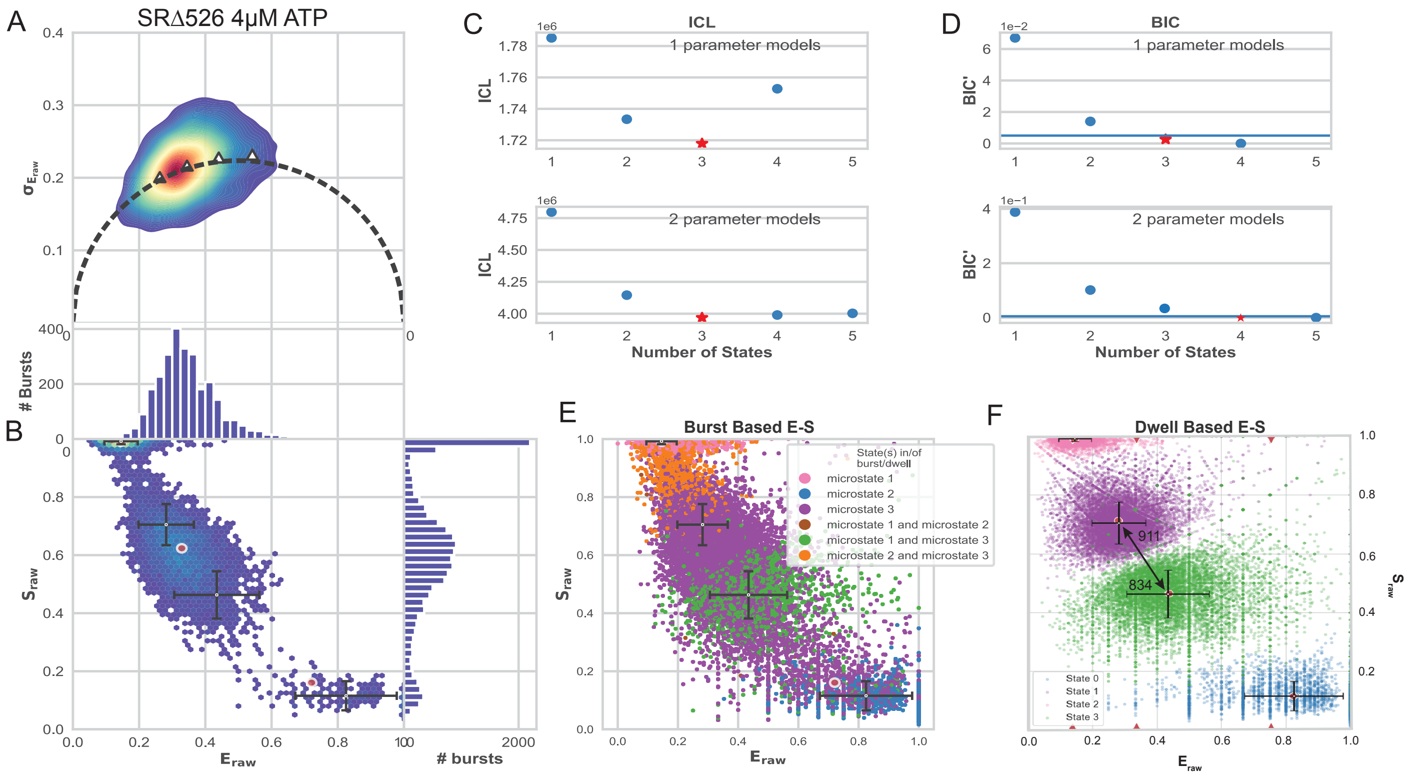


**Figure S6. Photon-by-photon H2MM analysis of SRΔ526 plus 4 μM ATP.**
(A) smFRET burst variance analysis (BVA) of 2762 bursts for SRΔ526 in the presence of 4 μM ATP identified by an all channel burst search (ACBS) (83, 84). (B) Stoichiometry (S) versus FRET efficiency (E) plot of raw smFRET burst data. H2MM analysis was performed using spH2MM (C and D, upper) and mpH2MM (C and D, lower) algorithms (67). Model performance as a function of increasing complexity was evaluated using (C) integrated complete likelihood (ICL) and (D) Bayesian information criterion (BIC) methods. The ‘star’ in each plot highlights the minimum number of states identified by each analysis. (E) Burst-based and (F) dwell-based plots of smFRET bursts colored by identified micro-state. The centroid of each micro-state distribution is shown (red and white dots), with error bars representing the standard deviation of the S and E values from Viterbi analysis. In the presence of 4 μM ATP, SRΔ526 is well described by two FRET micro-states, in addition to a donor-dark (D dark) and an acceptor-dark (A dark) state. Interconversion of the FRET micro-states, indicated by the double-headed arrow and observed rate constants ($s^{-1}$) in (F), occurs on the 1 msec time scale. For presentation clarity the interconversion pathways into and out of the D-dark and A-dark states are not shown.
